# Microglia determine ß-amyloid plaque burden but are non-essential for downstream pathology

**DOI:** 10.1101/2024.08.06.606795

**Authors:** Mosi Li, Kris Holt, Katherine Ridley, Jing Qiu, Kirsty Haddow, Deepali Vasoya, Xin He, Jane Tulloch, Declan King, David A Hume, Clare Pridans, Owen R Dando, Tara L Spires-Jones, Giles E Hardingham

## Abstract

Evidence points to a role for microglia in Alzheimer’s disease (AD) risk, although their position in the pathological cascade is incompletely understood, prompting us to generate a model of ß-amyloidopathy lacking microglia. We find evidence that microglia promote plaque formation and creation of an Aß fibril-rich zone surrounding the plaque core. However, plaque-proximal reactive astrogliosis, synapse loss, and neurite dystrophy are still observed in the absence of microglia.

## Results and Discussion

The APP_Swe_/PS1dE9 (APP/PS1) mice ^1^ (APP/PS1) ß-amyloidopathy mouse model exhibits extracellular Aß deposition, particularly in the neocortex, beginning at around 6 months ^2,3^. Reactive astrogliosis occurs proximal to plaques ^3,4^, as does synapse loss ^5^. We wanted to cross APP_Swe_/PS1dE9 mice onto the microglia-deficient Csf1r^ΔFIRE/ΔFIRE^ (FIRE^KO^) mouse line to assess whether Aß plaque deposition and downstream events are altered in brains lacking microglia. Of note however, the original FIRE^KO^ line was on a mixed background of CBA and C57BL/6J, and adverse phenotypes have been reported when crossed onto mice of C57BL/6J background, which happens to also be the background of the APP/PS1 mouse. When on a pure C57BL/6J background, the majority of FIRE^KO^ mice can succumb to hydrocephalus (Hume et al., in submission). Thus, prior to performing the genetic cross, both APP/PS1 and FIRE^KO^ lines were back-crossed several generations onto the CBA background. The four genotypes (± APP/PS1 ± FIRE^KO^) are hereafter referred to as WT-WT, WT-FIRE, APP-WT, APP-FIRE and all were 95-97% CBA background (Supplemental Fig. 1A). We confirmed an absence of microglia in FIRE and APP-FIRE mice by flow cytometry (Supplemental Fig. 1B, C) and IBA1 immunohistochemistry at the age studied (10-12 months, Supplemental Fig 1D), as well as by RNA-seq of neocortices followed by curation of microglia-enriched genes (Supplemental Fig. 1E). We also confirmed no differences (APP-WT vs. APP-FIRE) in mutant human APP expression (Supplemental Fig. 1F). Of note, the Swedish mutation (KM-NL) in the transgene increases the amyloidogenic processing of APP (initiated by ß secretase) to generate the 99 amino acid C-terminal fragment (as opposed to the non-amyloidogenic α-secretase processing to generate the 83 amino-acid CTF). We therefore measured the ratio of APP C-terminal fragments (99aa:83aam) As expected, the 99:83 ratio was found to be higher in APP-WT and APP-FIRE compared to WT-WT and WT-FIRE genotypes (Supplemental Fig. 1G). Also as expected, no difference was found between APP-WT and APP-FIRE genotypes (Supplemental Fig. 1G). We also confirmed that the humanized mutant APP and human PSEN1 transgenes were expressed at similar levels in APP-WT and APP-FIRE mice (Supplemental Fig. 1H,I).

To study Aß deposition we performed immunohistochemistry using three detection probes: i) the anti-Aß antibody 6E10; ii) the plaque core ß-sheet probe Methoxy-04 (MX-04); and iii) the anti-fibrillar Aß antibody OC. Compared to APP-WT mice, in APP-FIRE mice we observed a reduction in plaque number and overall burden in the neocortex in each of the three detection methods (Fig. 1A-C). Consistent with this reduction in aggregate burden, we saw a reduction in Aß found in SDS-insoluble, urea extractable brain extracts (Fig. 1D). We conclude from this, that microglia play an important role in Aß plaque formation.

**Figure 1.**
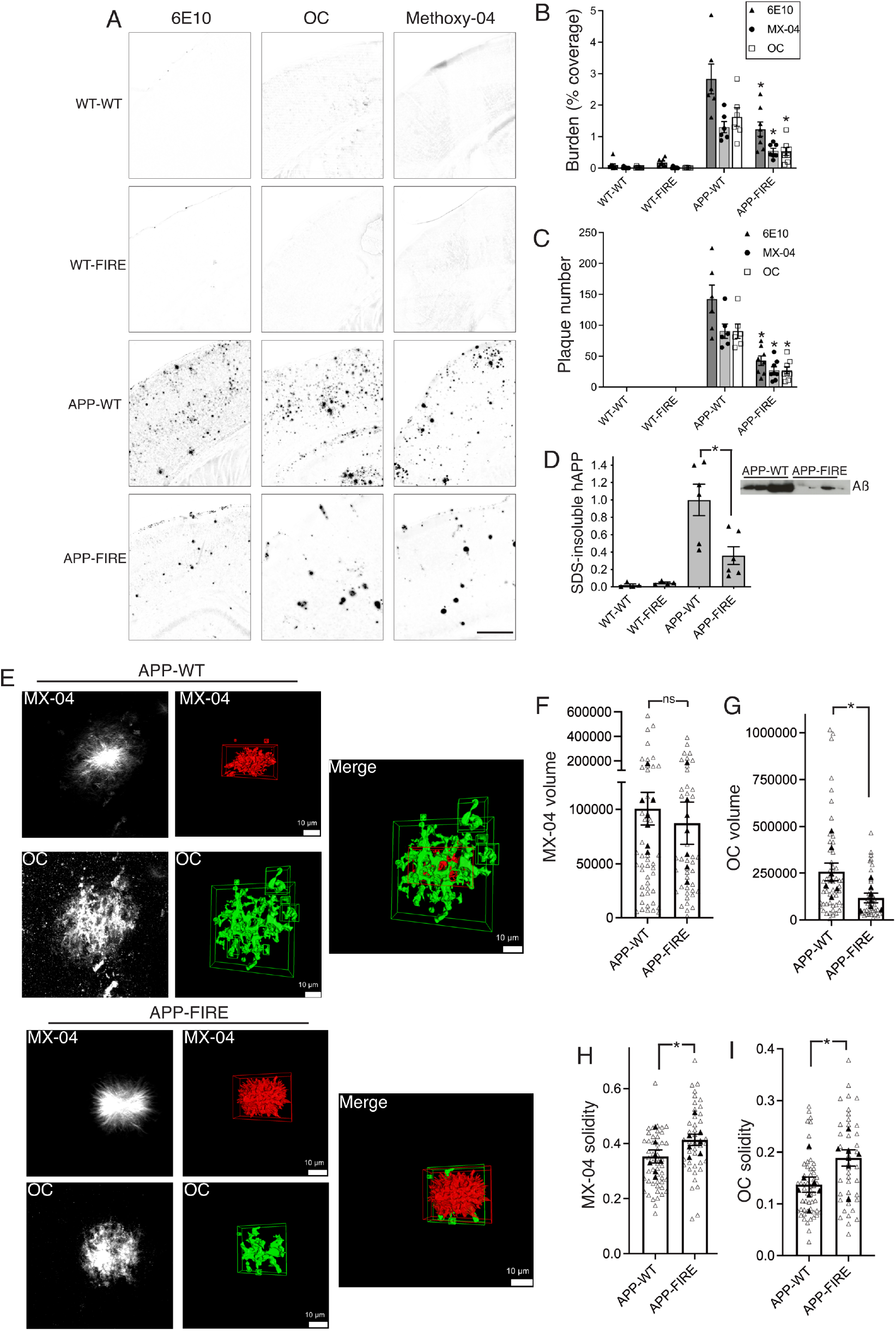
Absence of microglia causes a lowering of plaque burden and altered plaque properties. **A-C)** Analysis of neocortical Aß plaque burden in mice of the indicated genotypes (at 10-12 months) assessed by immunohistochemistry using antibodies against human APP (6E10), fibrillar Aß (OC) and the ß-sheet probe Methoxy-04. (A) shows example immunofluorescence images, inverted for easier visualisation (scale bar: 500 µm). (B) shows quantitation of coverage as a % of total area: F (3, 80)=87.35, p<0.0001 (2-way ANOVA main genotype effect), *p<0.0001 (APP-WT vs. APP-FIRE, Tukey’s post=hoc test). (C) shows analysis of plaque number. F (1, 36)=69.47, p<0.0001 (2-way ANOVA main genotype effect, APP-WT vs. APP-FIRE), *p<0.0001, 0.008, 0.008 (left-to-right, APP-WT vs. APP-FIRE, Tukey’s post=hoc test). N=10 mice (WT-WT), 7 (WT-FIRE), 6 (APP-WT), 8 (APP-FIRE). **E-I)** Sections of APP-WT and APP-FIRE mice (10-12 months) were subject to double immunohistochemistry with ß-sheet probe Methoxy-04 and anti-fibrillar antibody OC, and confocal z-stacks imaged. (E) shows example images in greyscale (maximum projection, left), 3D rendered images (middle), and combined 3D images (right). (F) shows quantitation of Methoxy-04 positive volume; (G) shows quantitation of OC-positive volume; (H) shows quantitation of the solidity of the Methoxy-04 positive volume; (I) shows quantitation of the solidity of Methoxy-04 volume. In all graphs, open triangles show individual plaque data points, filled triangles show the mean for each animal studied. F: “ns”: p=0.706 (t=0.386, df=12); G: p=0.030 (t=2.46, df=12); H: p=0.294 (t=2.472, df=12); I: p=0.033 (t=2.407, df=12), nested t-test treating plaques from the same animal as technical replicates, with different animals treated as biological replicates, n= 7 mice (APP-WT), 7 mice (APP-FIRE).

Although we observe a large effect of microglial absence on plaque deposition, sufficient numbers of plaques form to enable us to investigate plaque properties in the presence or absence of microglia. We studied individual plaques double-labelled with MX-04 and anti-OC. MX-04 preferentially labels the sense ß-sheet core of plaques, while the anti-fibrillar Aß antibody OC tends to label peri-core regions which are rich in fibrillar Aß (Fig. 1E). We observed that plaques in APP-WT and APP-FIRE mice had a similar volume of MX-04 immunoreactivity (Fig. 1F). However, plaques from APP-FIRE mice had slightly greater MX-04 solidity (Fig. 1H), defined as the proportion of pixels occupied by a bounding cuboid that occupies the entire shape. Analysis of the fibrillar component of plaques revealed that the anti-OC volume was lower in APP-FIRE mice compared to APP-WT mice (Fig. 1G) and had increased solidity (Fig. 1I). Moreover, the volume of OC staining that was negative for Methoxy-04 was also lower in in APP-FIRE mice compared to APP-WT mice (Supplemental Fig. 2A, B). Thus, in the absence of microglia, plaques have similar dense cores but possess a smaller volume of peri-core fibrillar Aß.

To further support these observations, we employed a supervised machine learning approach to predict microglial presence based on plaque morphology double-labelled with Methoxy-O4 and anti-fibrillar Aß OC antibody (Supplemental Fig. 2). The decision tree achieved high predictive power with 92% accuracy in distinguishing plaques from microglial-present and microglial-absent conditions (Supplemental Fig. 2C). Notably, the Methoxy-04:OC volume ratio, OC solidity, and OC major axis length emerged as key predictive features. This robust computational analysis supports our manual observations, indicating that changes in fibrillar amyloid properties are particularly indicative of microglial involvement in plaque formation and composition.

We next studied events downstream of Aß plaque deposition and their dependency on microglia, beginning with reactive astrogliosis. We focussed on reactive astrocyte marker GFAP and performed triple staining for plaque (6E10), reactive astrocytes (GFAP) and a 3^rd^ marker, ALDH1L1, which labels all astrocytes regardless of their reactive status. We measured the proportion of astrocytes (ALDH1L1-positive) that were reactive (GFAP-positive) in fields where a plaque was “present” (< 100 µm away), as well as “absent” (< 100 µm away in APP-WT and APP-FIRE mice, and all fields in WT-WT and WT-FIRE mice). In both APP-WT and APP-FIRE mice the proportion of astrocytes that were reactive was higher in plaque-present fields of view compared with plaque-absent areas, where reactivity was similar to the non-amyloidopathy genotypes WT-WT and WT-FIRE (Fig. 2A, B). However, the proportion of reactive astrocytes in plaque-containing fields of view was higher in APP-WT mice than APP-FIRE mice (Fig. 2B). Focussing on events < 25 µm from the plaque, we observed no difference in the proportion of reactive astrocytes (relative to total astrocytes) in APP-WT vs APP-FIRE mice. However, slightly further away (25-100 µm), the proportion of astrocytes that were reactive was higher in APP-WT vs APP-FIRE mice. Thus, there is no absolute requirement for microglia for plaque-proximal reactive astrogliosis to take place, although microglial presence causes greater levels of reactivity at intermediate distances from the plaque edge.

**Figure 2.**
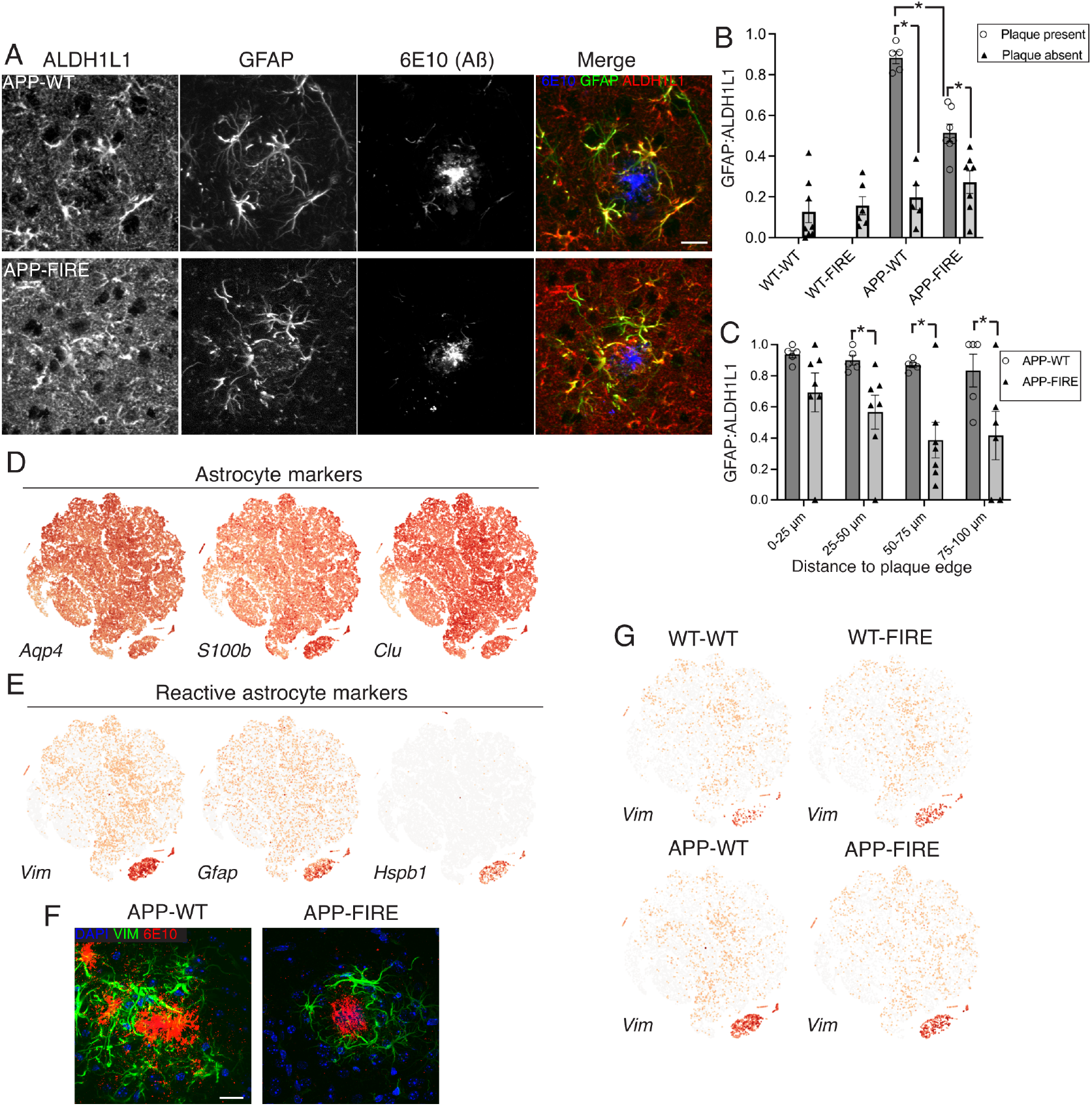
Aß-induced reactive astrogliosis has a microglia-independent component. **A-C)** Brain sections of mice of the indicated genotypes (at 10-12 months) were subject to triple immunohistochemistry with antibodies against reactive astrocytes (GFAP), all astrocytes (ALDH1L1), and Aß (OC). (A) shows example images (scale bar: 20 µm). (B) Within the neocortex the fraction of ALDH1L1-positive astrocytes that are also positive for GFAP was measured. “Plaque absent” refers to either non-APP genotypes or fields of view where no plaque is present and distance to the nearest plaque is > 100 µm. “Plaque present” refers to fields of view containing a plaque. Considering APP-WT and APP-FIRE genotypes, F (1,20)=19.42, p=0.0003, 2-way ANOVA main interaction effect of genotype and plaque presence. *p<0.0001, 0.0003, 0.0081 (left to right), n=5 mice (APP-WT), 7 mice (APP-FIRE). (C) Within fields of view containing a plaque, the fraction of ALDH1L1-positive astrocytes that are also positive for GFAP was measured and grouped according to distance to the edge of the plaque. F (1,39)=22.683, p<0.0001 (2-way ANOVA, main genotype effect), p=0.0366, 0.0033, 0.0124 (Sidak’s post-hoc test), n=5 mice (APP-WT), 7 mice (APP-FIRE). **D, E)** Astrocytes were sorted from neocortices of mice of all 4 genotypes (at 10-12 months) by FACS (ASCA-2 positive, O4-negative) and subject to single cell RNA-seq (10X Genomics). Minor contaminating cell types were removed and cells are shown as t-SNE projections. Expression of the indicated genes is shown (red: high expression; orange/yellow: low expression; grey: no expression) with example pan-astrocyte markers in (D) and reactive astrocyte markers in (E). 39,562 astrocytes are shown from the neocortices of n=4 mice (WT-WT), 4 (WT-FIRE), 4 (APP-WT), 3 (APP-FIRE). **F)** Example images showing vimentin expression in plaque-proximal astrocytes (scale bar: 20 µm). **G)** Data from the single cell RNA-seq expression were split according to genotype and expression of the reactive gene *Vim* shown.

We next wanted to know whether the transcriptional profile of reactive astrocytes was different in the absence of microglia. We performed single cell RNA-seq on astrocytes isolated by FACS from mice of all 4 genotypes, and confirmed the presence of markers *Aqp4, S100b* and *Clu* (Fig. 2D). We observed a cluster of astrocytes showing *Gfap* expression, as well as reactive markers *Vim* and *Hspb1* (Fig. 2E). We confirmed by immunohistochemistry against vimentin that these were the plaque-proximal astrocytes (Fig. 2F). These reactive astrocytes clustered together across the genotypes, suggesting that microglia do not strongly influence the transcriptional signature of reactive astrocytes proximal to Aß plaques (Fig. 2G). Note that small numbers of reactive astrocytes were observed in the non-APP/PS1 genotypes, (WT-WT and WT-FIRE), consistent with an age-dependent increase in basal reactive astrogliosis (Fig. 2G).

Since synapse loss is a major cause of cognitive decline in AD we wanted to study the impact of microglial deficiency in this process. We employed array tomography, employing pre- and post-synaptic markers (synaptophysin and PSD-95), scoring synapses where pre- and post-puncta pairs were within 50 nm of each other. We combined this with Methoxy-X04 to locate the plaques (Fig. 3A, B). We measured synapse density in APP-WT and APP-FIRE mice in the neocortex in fields of view containing plaques at their centre, compared to plaque-free regions, as well as WT-FIRE and WT-WT genotypes. We observed significant synapse loss in plaque-containing fields of view in both APP-WT mice and APP-FIRE mice, showing that microglia are not an absolute requirement for Aß-induced synapse loss (Fig. 3A-C). We then looked in more detail at plaque-containing fields of view, measuring synapse density as a function of distance to plaque core surface in 10 µm bins. We observed slightly less synapse loss in APP-FIRE mice up to 20 µm from the plaque surface, whereupon genotype differences were lost (Fig. 3D). We also quantified neurite dystrophies, defined by immunoreactivity to the marker SMI312 (an anti-phospho-neurofilament antibody) and a characteristic shape profile (see methods). Neurite dystrophy numbers were higher in plaque containing fields of view in both APP-WT mice and APP-FIRE mice, compared to plaque-free areas and plaque-free genotypes (Fig. 3E, F). However, the density of dystrophies was higher in APP-WT compared to APP-FIRE mice (Fig. 3F). Thus, plaque-associated synapse loss and neurite dystrophy do not have an absolute requirement for microglia, but both are quantitatively reduced when microglia are absent, potentially due to plaques possessing less fibrillar Aß around their edge.

**Figure 3.**
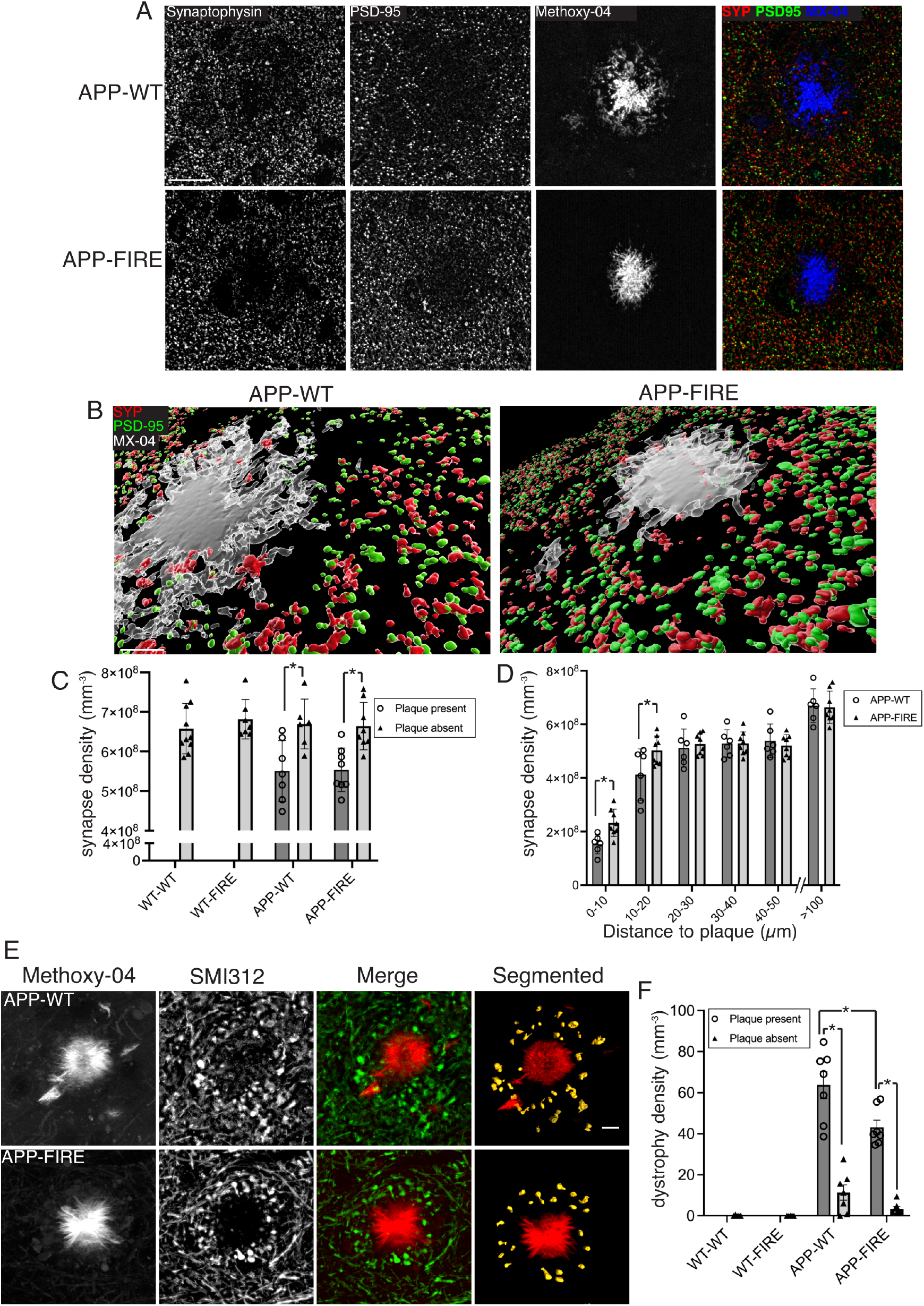
Microglial absence does not abolish plaque-associated synapse loss and neurite-dystrophy. **A)** Example pictures of single sections taken from mice at the indicated genotypes (10-12 months) that have been triple-labelled with plaque probe Methoxy-04, and pre-(SYP) and post-(PSD95) synaptic markers, as well as a psuedocoloured merged image on the right. Scale bar: 20 µm. **B)** Example 3D reconstruction of array tomography z-stacks. Scale bar: 5 µm. **C)** Synapse density (number per mm^3^) measured by array tomography in cortical sections of mouse brains of the indicated genotypes. “Plaque absent” refers to either non-APP genotypes or fields of view where no plaque is present and distance to the nearest plaque is > 100 µm. “Plaque present” refers to fields of view containing a plaque. F (1, 25)=23.10, p<0.0001, 2-way ANOVA (main effect of plaque presence, taking APP-WT and APP-FIRE data sets), p=0.0052, 0.0038 (Sidak’s post-hoc test). **D)** Within plaque-containing fields of view, synapse density was measured according to distance to the edge of a plaque. F (1, 60)=6.222, p=0.0154, (2-way ANOVA, main effect of genotype). *p=, 0.0489, 0.0193 (Sidak’s post-hoc test). N=10 mice (WT-WT), 7 (WT-FIRE), 6 (APP-WT), 8 (APP-FIRE). **E, F)** APP-WT and APP-FIRE brain sections were subject to double immunohistochemistry to assess plaque-proximal neurite dystrophy using the Aß plaque probe Methoxy-04 and the anti phospho-neurofilament antibody SMI312, with SMI312 reactive dystrophies defined by morphological criteria (see Methods). Scale bar: 10 µm. In (F) the density of neurite dystrophies (per µm^3^) is assessed in plaque-containing fields of view (“plaque present”) and regions lacking plaques (“plaque absent”) of APP-WT and APP-FIRE mice, and a lack of dystrophies confirmed in the non-plaque-containing genotypes (WT-WT and WT-FIRE). F(1,24)=120.7, p<0.0001 (2-way ANOVA main effect of plaque proximity), p=0.0024 (2-way ANOVA main effect of genotype). *P<0.0001, 0.004, <0.0001 (Sidak’s post-hoc test). N=5 mice (WT-WT), 5 (WT-FIRE), 7 (APP-WT), 7 (APP-FIRE).

The impact of microglial absence on plaque burden suggest that the overall consequences of mutant APP and PSEN1 expression for the brain may be diminished when microglia are absent. To study this further we looked at the impact of the APP_Swe_/PS1dE9 on neocortical gene expression by RNA-seq both against a wild-type background (APP-WT vs. WT-WT) as well as a FIRE^KO^ background (APP-FIRE vs. WT-FIRE). We detected 813 protein-coding genes whose expression was significantly altered in the APP-WT mouse, relative to WT-WT mice (>1 FPKM, p_adj<0.05, Supplemental Fig. 3A). GO analysis revealed an inflammatory/immune signature in the genes induced, as well as a cellular response to Aß, as well as numerous other pathways in the up- and down-regulated sets (Supplemental Fig. 3B, C). In contrast, only 22 genes were significantly altered in the APP-FIRE mouse compared to WT-FIRE (Supplemental Fig. 3D). Since microglia are a key contributor to neuroinflammation, the loss of the inflammatory/immune signature is expected. However, the near-abolition of all transcriptional changes is likely to also be a consequence of lower burden of plaques and peri-plaque lower-order polymers in APP-FIRE mice.

A previous study aimed to investigate the impact of microglial deficiency on ß-amyloidopathy by crossing the FIRE^KO^ mouse onto the 5xFAD ß-amyloidopathy model ^6^. However, this cross resulted in severe cerebrovascular amyloid angiopathy, haemorrhage and premature lethality, commencing at around 1 month and rising to 70% death by 6 months. A combination of the poor health of the FIRE^KO^ allele on the C57BL6/J background, plus the aggressive nature of the amyloidopathy model, may be contributing factors to the lethality reported. Regardless of the mechanism, the underlying progressive and unpredictable early lethality observed by Shabestari et al. makes understanding the role of microglia in pre- and post-plaque cellular processes difficult, something we have overcome by crossing our lines onto the CBA background and utilizing a slowly progressing model of ß-amyloidopathy. Of note, we observed no difference in the quantity of vascular Aß between of APP-WT and APP-FIRE mice, measured by colocalization of Aß with collagen IV, (Supplemental Fig. 4A, B). Our study revealed that genetic loss of microglia causes a dramatic decrease in plaque load.

Others have studied the impact of killing microglia, with mixed results. The CSF1R antagonist PLX5622 leads to effective (but not 100% efficient) microglial death (and has off-target effects ^7^) and was reported to reduce plaque load in the fast-progressing 5xFAD mouse ^8^. In contrast, a different strategy to kill microglia had no effect on plaque burden in the APP/PS1 mouse ^9^. The FIRE^KO^mouse has the advantage over an active depletion strategy in that the brain is not left with a substantial pool of dead cells. Moreover, the loss of microglia in the FIRE^KO^mouse is essentially total, meaning that the degree of microglia-independent plaque deposition can be accurately defined, unlike in previous studies where plaque formation was attributed to areas of incomplete microglial death ^8^.

Our study raises the question as to why microglial absence reduces plaque load, a finding which is contrary to the classical view of microglia as involved in Aß clearance and inhibition of plaque seeding ^10^. It could be that microglial phagocytosis of Aß provides conditions conducive to aggregation. Aß aggregation has been proposed to occur more efficiently when taken up into the endosomal/lysosomal system due to the lower pH and higher local concentration ^11^. Interestingly, an imaging study demonstrated that plaque-proximal microglia can phagocytose Aß and release it upon cell death, contributing to plaque growth ^12^. Recently, a study proposed that microglia play a role in Aß plaque dense core formation via TAM receptor signaling ^13^. While this offers a potential explanation for the effect of microglial absence in our study, it is notable that in APP-FIRE mice, plaques have a similar Methoxy-04 reactive dense core as in APP-WT mice (Fig. 1F), albeit in reduced numbers. This points to the possibility that plaque seeding and plaque core growth are separate processes, and that the former has a stronger dependency on microglia rather than the latter.

In the context of synapse loss in AD, studies in models of ß-amyloidopathy have proposed microglial phagocytosis as a mechanism, involving complement components or other synaptic “eat me” signals like phosphatidylserine ^14-16^. However, we observe significant Aß plaque-proximal synapse loss in the absence of microglia (and by extension, an absence of C1q, Supplemental Fig. 1E). The very small impact of microglial absence on plaque-proximal could be due to the loss of microglia-mediated synapse loss, but equally could be due to the reduced halo of lower-order Aß species around the plaque core in APP-FIRE mice (Supplemental Fig 2A, B). Regardless, the substantial level of microglia-independent synapse loss points to other, potentially neuron-intrinsic mechanisms contributing. For example, Aß has been reported to induce excitotoxic synapse loss in an NMDA receptor dependent manner ^17-19^. The altered function of peri-plaque astrocytes may also be a contributing factor to synapse loss ^20^. Indeed, the fact that Aß-induced reactive astrocytes exhibit hypo-expression of glutamate transporters may cause glutamate dyshomeostasis in this region, increasing excito-toxicity.

In summary, we have placed microglia at a critical early stage of AD pathology, consistent with strong genetic evidence for their role in AD initiation, as well as defining their limited contribution to synapse loss downstream of plaque formation. Given this, microglial-centric genetic risk may functionally impact on microglial capacity to seed amyloid plaques, and targeting microglia for successful treatment of AD may require the patient to be at an early stage of disease.

## Acknowledgements

This work was funded by the UK Dementia Research Institute which receives its funding from UK DRI Ltd, core funded by the UK Medical Research Council. We thank Matthew Nolan for access to slide scanning facilities, Andrea Corsinotti and Meryam Beniazza at the Edinburgh FACS and single cell RNA-seq facility, and Paul Baxter for assistance in confocal imaging.

## Figures

**Supplemental Figure 1.**
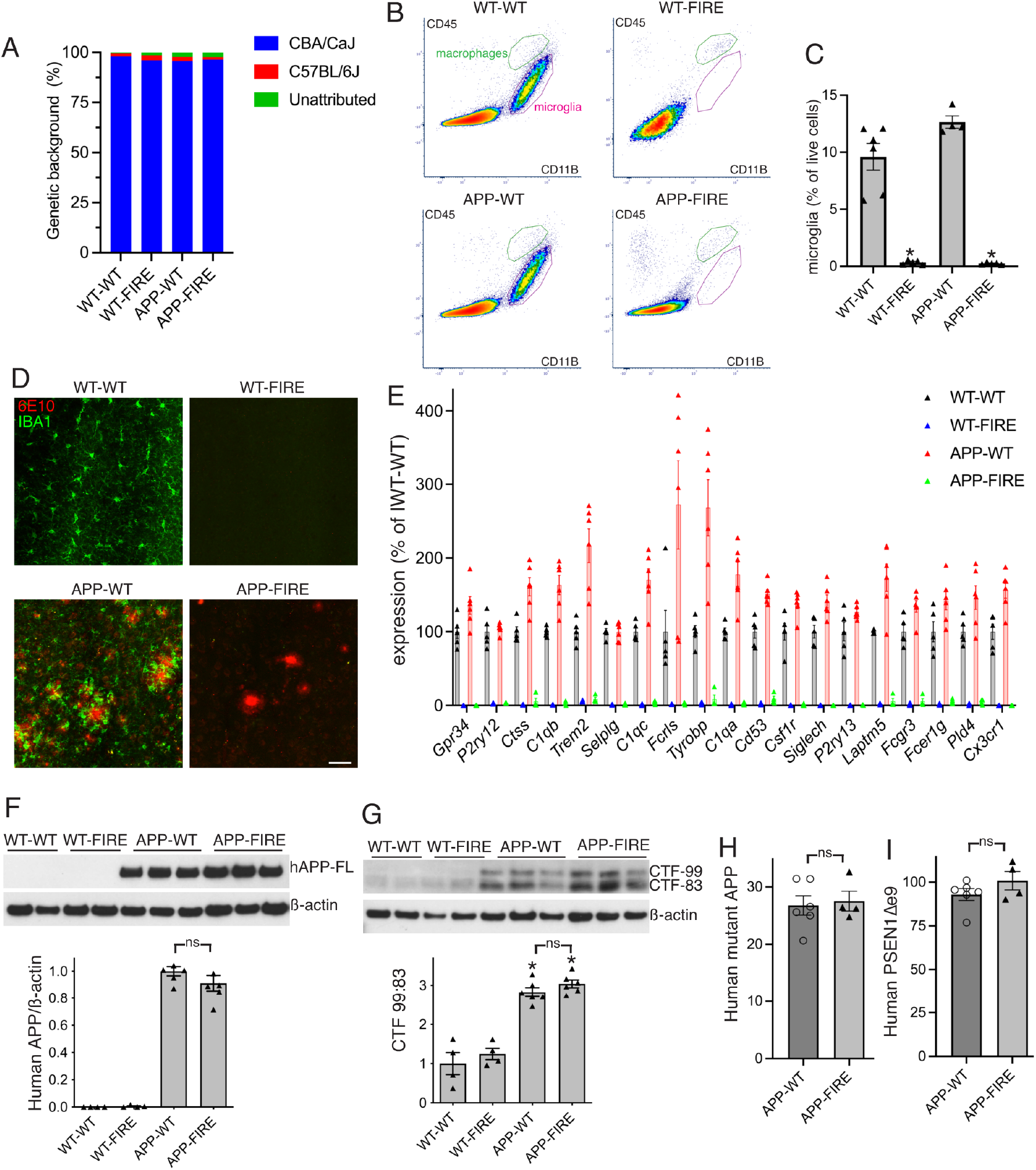
**A)** Genetic background of the indicated genotypes, assessed using the mini mouse universal genotyping array ^21^ performed by Transnetyx, is shown. **B, C)** Flow cytometry of neocortices of the indicated genotypes (aged 10-12 months), showing just cells with high Cd11b expression, plotted against Cd45 expression. Approximate region comprised of microglia is shaded in green, with border associated macrophages (High Cd45-Cd11b ratio) shown in white. (C) shows quantitation, with individual data points relating to individual animals. *P<0.0001, one-way ANOVA followed by Tukey’s post-hoc test. **D)** Example immunofluorescence images of the indicated genotypes, with 6E10-reactive plaques and IBA-1 positive microglia shown. Scale bar: 50 µm. **E)** RNA sequencing data showing the relative expression of microglia-specific genes in neocortical tissue of the indicated genotypes at 10-12 months. Genes were selected based on previously published single-cell sequencing data ^22^ that described 265 subclusters in the CNS and PNS. To define microglia-enriched genes, we calculated the fold-enrichment of expression in the “MGL1” subcluster relative to the next highest expressing non-immune cell subcluster, requiring an average of ≥1 FPKM in wild-type mice. Data relating to genes with a microglial enrichment of ≥50-fold are shown. Individual data points relate to individual animals. N=5 mice (WT-WT), 6 (WT-FIRE), 6 (APP-WT), 4 (APP-FIRE). Relative to WT-WT, all genes in WT-FIRE and APP-FIRE mice are significantly down-regulated (DESeq2 adjusted p-value <0.05) **F)** Expression of human full-length APP in neocortices (10-12 months) assessed by western blot with the anti-(human) APP antibody 6E10, normalised to ß-actin. *p<0.0001, relative to WT-WT, one-way ANOVA followed by Tukey’s post-hoc test. “ns”: p=0.268. N=4 mice (WT-WT), 4 (WT-FIRE), 6 (APP-WT), 6 (APP-FIRE). **G)** Assessment of the relative expression of 99-amino acid vs. 83-amino acid C-terminal APP fragments, normalized to the average WT-WT level, assessed using an antibody against the APP C-terminus (C1/6.1). *p<0.0001, relative to WT-WT, one-way ANOVA followed by Tukey’s post-hoc test. ns: p=0.714. N=4 mice (WT-WT), 4 (WT-FIRE), 6 (APP-WT), 6 (APP-FIRE). **H, I)** RNA sequencing data showing the expression of human mutant transgenes *APP* (H) and *PSEN1* (I) in the indicated genotypes in the neocortex (10-12 months) expressed as number of human-specific APP reads per million reads. Human-specific reads were identified using our Sargasso workflow ^23^, ns: p=0.770 (H), 0.218 (I), 2-tailed unpaired t-test. N=6 mice (APP-WT), 4 (APP-FIRE).

**Supplemental Figure 2.**
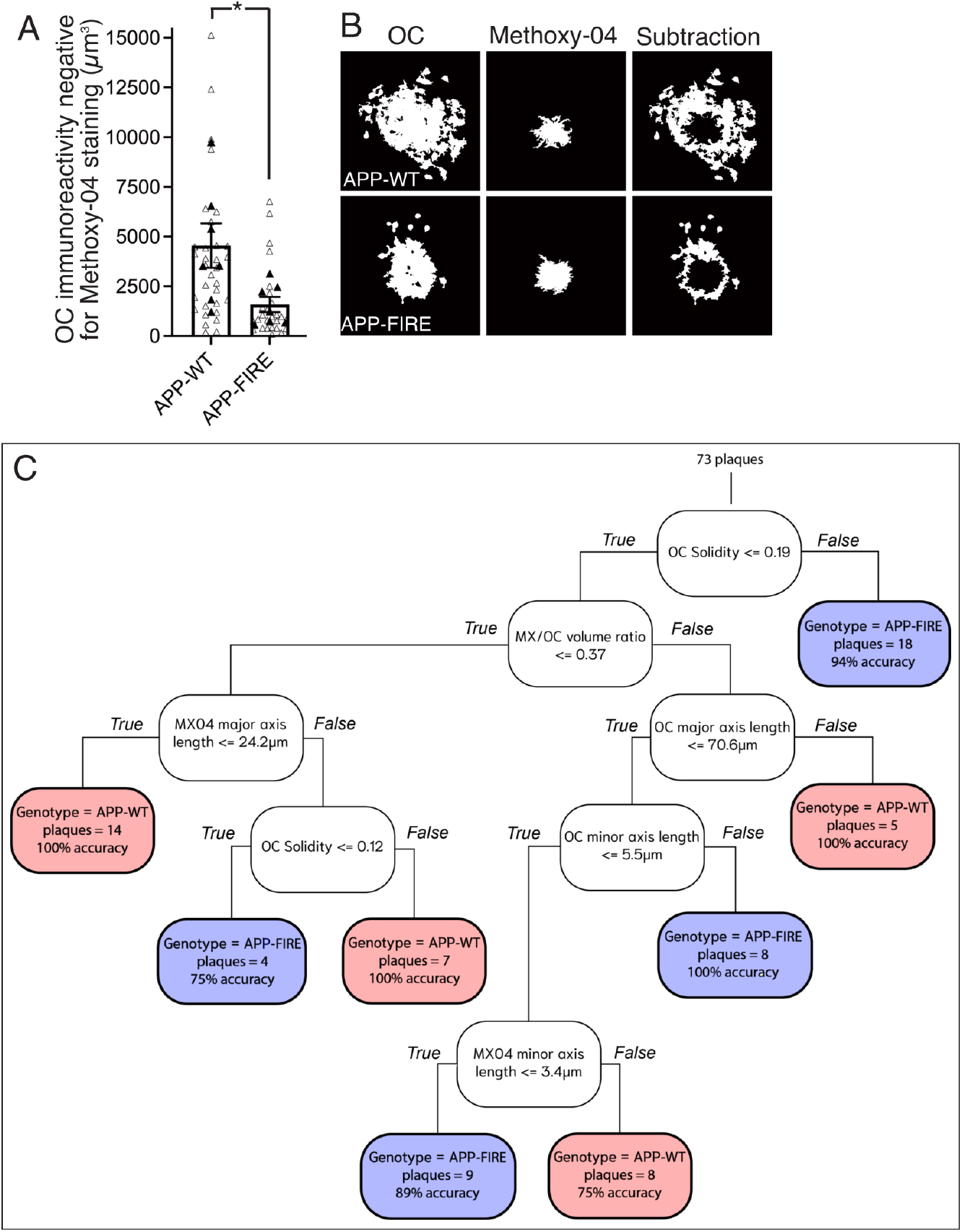
**A, B)** Quantitation of OC-positive volume that is negative for Methoxy-04 staining. Open triangles show individual plaque data points, filled triangles show the mean for each animal studied. *p=0.0297 (t=2.466, df=12), nested t-test, treating plaques from the same animal as technical replicates, with different animals treated as biological replicates, n= 7 mice (APP-WT), 7 mice (APP-FIRE). (B) shows example images. **C)** Representation of a machine learning-derived Decision Tree to predict mouse genotype based on images of Aß plaques (double labelled with Methoxy-O4 and anti-fibrillar Aß (OC antibody)). Analysis was made of 92 plaques from a total of 14 animals (7 APP-WT and 7 APP-FIRE). Grid search optimization with five-fold cross-validation was used to identify the optimal parameters for a decision tree classifier. This process yielded a model with entropy criterion, maximum depth of 8, minimum samples per leaf of 4, and minimum samples for split of 10. We then applied a 20:80 train:test split to evaluate the model’s performance. 73 plaques were tested as shown in the figure, with various decision points based on morphological properties leading to genotype prediction. Overall, the tree had predictive power of 92%.

**Supplemental Figure 3.**
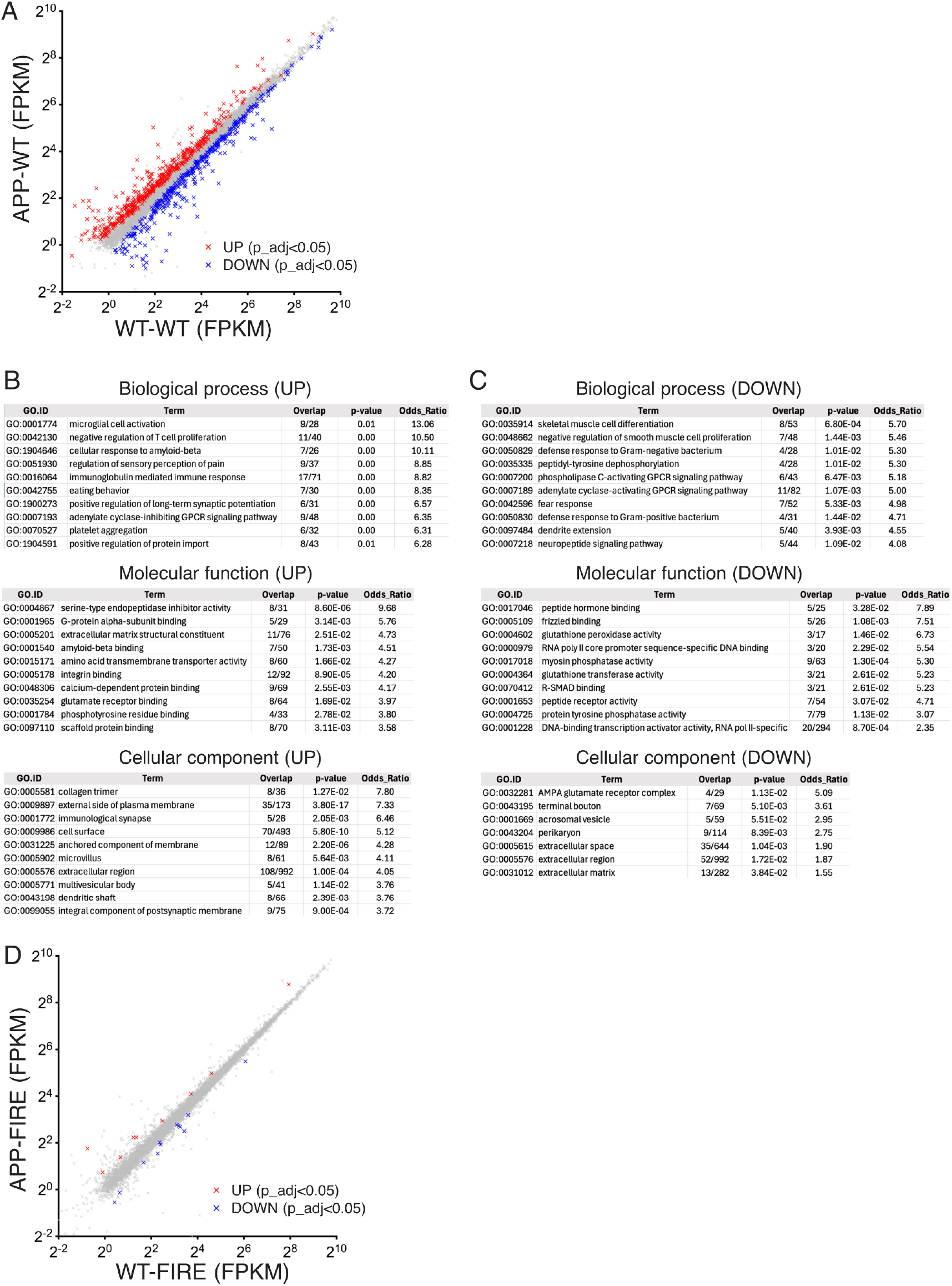
**A)** RNA-seq was performed on RNA extracted from cortical tissue taken from WT-WT (n=5) and APP-WT (n=6) mice (10-12 months). Expression levels for protein-coding genes (average expression >1 FPKM is shown, with the 813 significantly regulated genes (DESeq2 adjusted p<0.05) highlighted in red (induced) or blue (repressed). **B, C)** GO gene set enrichment analysis for genes up-regulated (B) or down-regulated (C) in the APP-WT cortex compared to WT-WT. Gene sets with ≥20 members in the background were considered, and the top 10 (by odds ratio) are shown (or fewer if 10 do not reach significance). **B)** RNA-seq was performed on RNA extracted from cortical tissue taken from WT-FIRE (n=6) and APP-FIRE (n=4) mice (10-12 months). Expression levels for protein-coding genes (average expression >1 FPKM is shown, with the 22 significantly regulated genes (DESeq2 adjusted p<0.05) highlighted in red (induced) or blue (repressed).

**Supplemental Figure 4.**
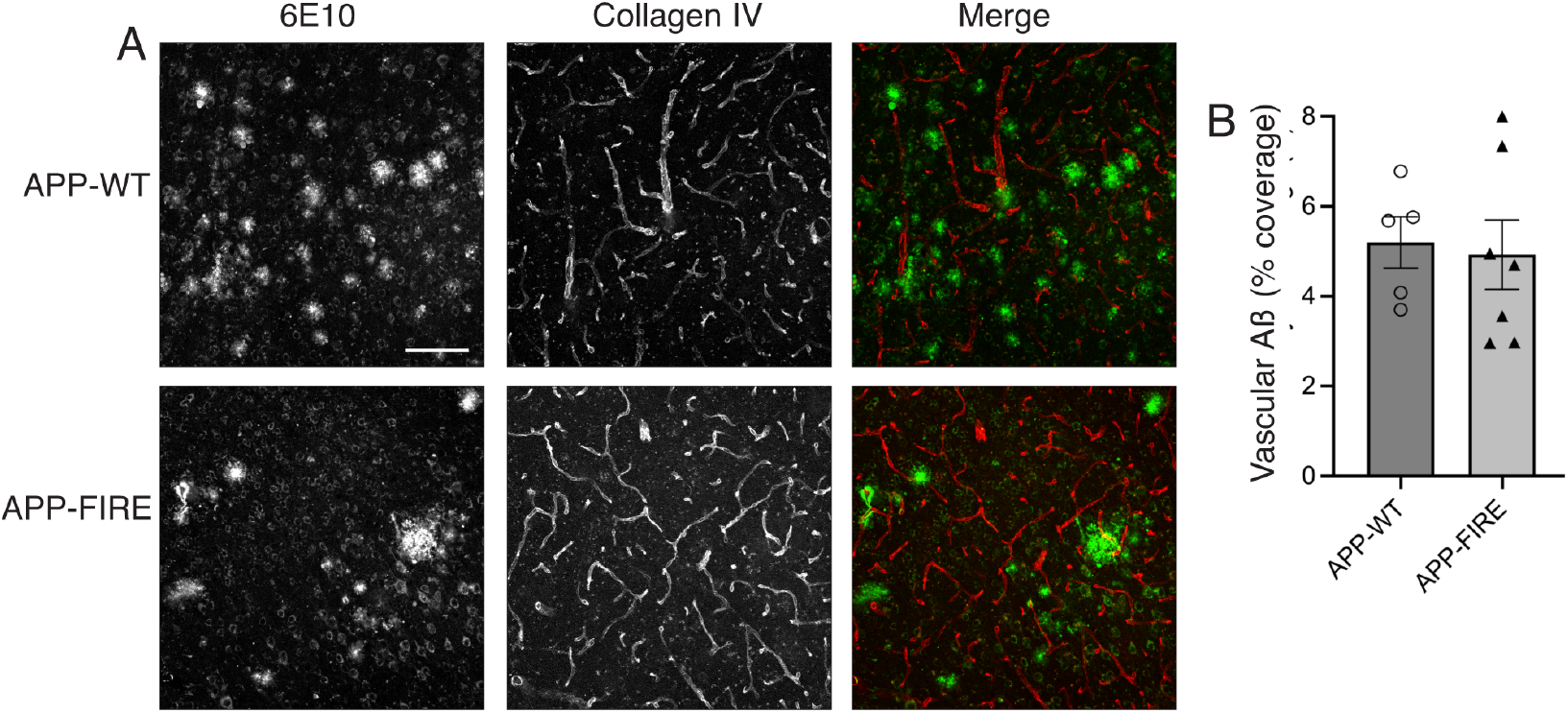
**A, B)** APP-WT and APP-FIRE brain slices (10-12 months) were subject to double immunohistochemistry using antibodies against human APP (6E10) and the vasculature-associated Collagen IV. The percentage of Collagen IV immunoreactivity which also co-localised with 6E10 immunoreactivity was measured and found to be similar in the two genotypes. N= 5 APP-WT mice, n=7 APP-FIRE mice.

## References

1 Jankowsky, J. L. et al. Mutant presenilins specifically elevate the levels of the 42 residue betaamyloid peptide in vivo: evidence for augmentation of a 42-specific gamma secretase. Human molecular genetics 13, 159–170 (2004). 10.1093/hmg/ddh019

2 Garcia-Alloza, M. et al. Characterization of amyloid deposition in the APPswe/PS1dE9 mouse model of Alzheimer disease. Neurobiol Dis 24, 516–524 (2006). 10.1016/j.nbd.2006.08.017

3 Jiwaji, Z. et al. Reactive astrocytes acquire neuroprotective as well as deleterious signatures in response to Tau and Aß pathology. Nat Commun 13, 135 (2022). 10.1038/s41467-021-27702-w

4 Kamphuis, W. et al. GFAP isoforms in adult mouse brain with a focus on neurogenic astrocytes and reactive astrogliosis in mouse models of Alzheimer disease. PLoS ONE 7, e42823 (2012). 10.1371/journal.pone.0042823

5 Koffie, R. M. et al. Oligomeric amyloid beta associates with postsynaptic densities and correlates with excitatory synapse loss near senile plaques. Proc Natl Acad Sci U S A 106, 4012–4017 (2009). 10.1073/pnas.08116981060811698106 [pii]

6 Kiani Shabestari, S. et al. Absence of microglia promotes diverse pathologies and early lethality in Alzheimer’s disease mice. Cell Rep 39, 110961 (2022). 10.1016/j.celrep.2022.110961

7 Lei, F. et al. CSF1R inhibition by a small-molecule inhibitor is not microglia specific; affecting hematopoiesis and the function of macrophages. Proc Natl Acad Sci U S A 117, 23336–23338 (2020). 10.1073/pnas.1922788117

8 Spangenberg, E. et al. Sustained microglial depletion with CSF1R inhibitor impairs parenchymal plaque development in an Alzheimer’s disease model. Nat Commun 10, 3758 (2019). 10.1038/s41467-019-11674-z

9 Grathwohl, S. A. et al. Formation and maintenance of Alzheimer’s disease beta-amyloid plaques in the absence of microglia. Nat Neurosci 12, 1361–1363 (2009). 10.1038/nn.2432

10 Long, J. M. & Holtzman, D. M. Alzheimer Disease: An Update on Pathobiology and Treatment Strategies. Cell 179, 312–339 (2019). 10.1016/j.cell.2019.09.001

11 Hu, X. et al. Amyloid seeds formed by cellular uptake, concentration, and aggregation of the amyloid-beta peptide. Proc Natl Acad Sci U S A 106, 20324–20329 (2009). 10.1073/pnas.0911281106

12 Baik, S. H., Kang, S., Son, S. M. & Mook-Jung, I. Microglia contributes to plaque growth by cell death due to uptake of amyloid beta in the brain of Alzheimer’s disease mouse model. Glia 64, 2274–2290 (2016). 10.1002/glia.23074

13 Huang, Y. et al. Microglia use TAM receptors to detect and engulf amyloid beta plaques. Nat Immunol 22, 586–594 (2021). 10.1038/s41590-021-00913-5

14 Hong, S. et al. Complement and microglia mediate early synapse loss in Alzheimer mouse models. Science 352, 712–716 (2016). 10.1126/science.aad8373

15 Salter, M. W. & Stevens, B. Microglia emerge as central players in brain disease. Nat Med 23, 1018–1027 (2017). 10.1038/nm.4397

16 Rueda-Carrasco, J. et al. Microglia-synapse engulfment via PtdSer-TREM2 ameliorates neuronal hyperactivity in Alzheimer’s disease models. EMBO J 42, e113246 (2023). 10.15252/embj.2022113246

17 Sharma, M., Pal, P. & Gupta, S. K. The neurotransmitter puzzle of Alzheimer’s: Dissecting mechanisms and exploring therapeutic horizons. Brain Res 1829, 148797 (2024). 10.1016/j.brainres.2024.148797

18 Ong, W. Y., Tanaka, K., Dawe, G. S., Ittner, L. M. & Farooqui, A. A. Slow excitotoxicity in Alzheimer’s disease. J Alzheimers Dis 35, 643–668 (2013). 10.3233/JAD-s121990

19 Tu, S., Okamoto, S., Lipton, S. A. & Xu, H. Oligomeric Abeta-induced synaptic dysfunction in Alzheimer’s disease. Mol Neurodegener 9, 48 (2014). 10.1186/1750-1326-9-48

20 Patani, R., Hardingham, G. E. & Liddelow, S. A. Functional roles of reactive astrocytes in neuroinflammation and neurodegeneration. Nat Rev Neurol 19, 395–409 (2023). 10.1038/s41582-023-00822-1

21 Sigmon, J. S. et al. Content and Performance of the MiniMUGA Genotyping Array: A New Tool To Improve Rigor and Reproducibility in Mouse Research. Genetics 216, 905–930 (2020). 10.1534/genetics.120.303596

22 Zeisel, A. et al. Molecular Architecture of the Mouse Nervous System. Cell 174, 999–1014 e1022 (2018). 10.1016/j.cell.2018.06.021

23 Qiu, J. et al. Mixed-species RNA-seq for elucidation of non-cell-autonomous control of gene transcription. Nat Protoc 13, 2176–2199 (2018). 10.1038/s41596-018-0029-2

