## Supplemental Methods for "Microglia determine ß-amyloid plaque burden but are non-essential for downstream pathology"

***Animals:*** All animal procedures were conducted in accordance with the UK Animal (Scientific Procedures) Act 1986 under the authority of project licence number PP8710936, with local ethical and veterinary approval at the University of Edinburgh. This study used Csf1r^∆FIRE/∆FIRE^ (FIRE KO) mice ^1^, which have been backcrossed for several generations onto CBA mice achieve a predominantly CBA background. These mice were then mated with APP^Swe^/PS1dE9 (APP/PS1) mice ^2^, which were similarly been backcrossed to a CBA background. The resulting experimental line possesses a CBA background greater than 95%. Both male and female mice aged 9-13 months were used throughout the study. The mice were group-housed in environmentally enriched cages within humidity- and temperature-controlled rooms, maintained on a 12-hour light-dark cycle with free access to food and water. Mouse genotypes were determined using genomic DNA extracted from ear biopsy tissue. Real-time PCR with transgene-specific probes and the primer sequences listed below was performed (Transnetyx, Cordova, TN), FIRE primers: Fwd: GCGGTTGTAGGAAACCCTGA; Rev: CTTCCAAGTGTCTCCTGCTG; APP/PS1 primers: Fwd: AGATGAGCCACGCAGTCC; Rev: TCACAGAAGATACCGAGACT.

***FACS and single cell RNA-seq:*** Cerebral neocortices were collected then dissociated with Adult Brain Dissociation Kit (Miltenyi Biotec) according to the manufacturer’s protocol on a gentleMACS Octo Dissociator using program 37C_ABDK_01. Dissociated samples were then treated with debris and red blood cell removal steps to obtain cell suspensions. For FACS sorting astrocytes, cells were resuspended in a final volume of 100 µl DPBS + 0.5% BSA buffer. Then cells were incubated with fluorescent conjugated antibodies at 4° C for 30 minutes. The following antibodies were used: CD 11b-FITC (1:500, Biolegend, #01206), ACSA2 APC (1:200, Miltenyi Biotec, #130-116-245), O4 PE (1:100, Miltenyi Biotec, #30-117-357), CD45-Pacific Blue (1:100. Biolegend, #103126) 1:100, LY6G Brilliant Violet 711 (1:100, Biolegend, #127643) 1:100. ACSA2 APC (+) and O4-PE (-) live single cells were sorted for single cell RNA sequencing experiment. FACS experiments were performed on a BD FACSAria II Cell Sorter (BD Biosciences).

Single cell astrocytes suspensions were loaded onto a 10x Chromium Controller (10x Genomics) according to the manufacturer’s protocol using 10x Genomics proprietary technology. Single cell transcriptomic amplification and library preparation were conducted with the Chromium Single Cell 3′ v3.1 Reagent Kit (10x Genomics), following the manufacturer’s instructions. After preparation, libraries underwent sequencing on the Sequencing was performed on the iSeq 100 System (Illumina Inc, #20021532) using the iSeq 100 i1 Reagent v2 (300 cycle) Kit (#20031371). Library molarity for sequencing was calculated using the Qubit dsDNA quantification results and the fragment size information from the Bioanalyser results. Libraries were normalised to 10nM and equal volumes were pooled and diluted for sequencing. PhiX Control v3 (#FC-110-3001) library was spiked into the run at a concentration of ~4% to help with cluster resolution and facilitate troubleshooting in case of any problems with the run. Data was returned to the Investigator to allow the calculation of the mean reads per cell for each sample so this information could be used to rebalance the pool for deeper sequencing. Sequencing was then performed on the NextSeq 2000 platform (Illumina Inc, #SY-415-1002) using the NextSeq 1000/2000 P3 Reagents (100 cycles) v3 Kit (#20040559). Normalised libraries previously run on the iSeq 100 were pooled at different volumes for each runs (2 in total) following the investigator’s pooling strategies. PhiX Control v3 (#FC-110-3001) library was spiked into both run at a concentration of ~1%. sequencing reads were mapped to the mouse genome, and per-cell, per-gene count matrices produced, using 10x CellRanger version 7.0.1 ^3^. QC, normalisation and clustering of data was performed using the Seurat R package, version 4.4.0 ^4^. Doublets were identified and removed using the ScDblFinder R package, version 1.10.1 ^5^. Pseudo-bulk differential expression analysis was performed by summarising single cell gene expression profiles at the subject level using the aggregateBioVar R package, version 1.6.0, then differentially expressed genes between genotypes were calculated using DESeq2.

***Bulk RNA-seq:*** RNA extraction was performed using the RNeasy Mini Kit (Qiagen, #74804) following the manufacturer's instructions. Briefly, tissue samples were homogenized in 1 mL of QIAzol lysis reagent. After adding 200 µL of chloroform and centrifugation, the aqueous phase was collected and mixed with an equal volume of ethanol. RNA was then extracted and purified using RNeasy Mini spin columns, involving several washing steps. Finally, the purified RNA was eluted in 30 µL of RNase-free water. RNA sequencing of neocortical samples was performed using TruSeq Stranded mRNA-seq library preparation (500 ng of starting material was used), followed by next-generation sequencing on the Illumina NovaSeq S1 Flowcell (100 cycles). samples were sequenced to a depth of approximately 45 million 100 base-pair, paired-end reads. Reads were mapped to the primary assembly of the mouse (GRCm39) reference genome contained in Ensembl release 110, using the STAR RNA-seq aligner, version 2.7.9a ^6^. Tables of per-gene read counts were generated from the mapped reads with featureCounts, version 2.0.2 ^7^, and differential gene expression was performed in R using DESeq2, version 1.36.0 ^8^.

***Immunohistochemistry:*** At the experimental endpoint, mice were intracardially perfused with phosphate-buffered saline (PBS) under terminal anaesthesia. The brains were then post-fixed in 4% paraformaldehyde (PFA) overnight, followed by immersion in 30% sucrose for 72 hours. Afterward, the brains were snap-frozen and stored at −80 °C. Coronal sections (50 μm) were prepared using a cryostat (Leica). The sections were permeabilized and blocked with 1% bovine serum albumin (BSA) and 0.5% Triton X-100 in PBS for 1 hour at room temperature (RT), followed by incubation with primary antibodies diluted in the blocking buffer at 4 °C overnight. Primary antibodies used: anti-hAPP 6E10 (#803001, 1:500, Biolegend), Anti-Aß fibril OC antibody (#AB2286, 1:500, Millipore), anti-ALDH1L1 (#702573, 1:400, ThermoFisher), anti-GFAP (#13-0300, 1:1000, ThermoFisher), Collagen Type IV (#AB769, 1:100, Millipore), anti-IBA1 (#1014660-1, 1:500, Abcam), anti-vimentin (#GR3429646-9, 1:500, Abcam), anti-neurofilament SMI312 (#SMI-312R, 1:500, Covance).

Following primary antibody incubation, tissue was washed a further 3 times and incubated with secondary antibodies (Table 2) for a period of 2 hours at room temperature, under agitation, while protected from light. For experiments involving additional post-staining with Methoxy-X04 (MX04), 10mg/ml MX04 stock in DMSO was diluted to a working concentration of 50 µM in kolliphor EL (Sigma-Aldrich; C5135-500G) added in a 2:1 volume ratio before topping up with 1X PBS. Tissue was then incubated in working MX04 solution for 15 minutes at room temperature under agitation with samples protected from light prior to a final 3 washes with 1X PBS. Sections were mounted on 25x75x1mm SuperFrostTM Adhesion slides (Epredia; J1800AMNZ) before sealing with 24 x 40 mm 100 pcs borosilicate coverslips. VectaShield Antifade mounting medium with DAPI (Vector Laboratories; l H-1200) was used for mounting, excepting experiments involving post-staining with MX04, in which case VectaShield Antifade mounting medium without DAPI (Vector Laboratories; H-1000) was used.

***Analysis of plaque burden:*** For whole brain analysis, whole hemibrains were imaged using a Zeiss Axio Scan Z1 slide scanner, fitted with a Hamamatsu Orca Flash camera (Zeiss, Baden-Württemberg, Germany). Images were acquired as maximum intensity projections of tile scanned z-stacks at 10X magnification and saved as 16-bit greyscale images in .czi format. For analysing tile-scanned maximum intensity projections, FIJI/ImageJ was used for all quantification by an experimenter blind to genotype. Channels from each raw .czi image (16-bit greyscale format) were split and saved in .tif format. Regions of interest (ROIs) were then drawn manually to delineate the cerebral cortex and hippocampus. For the cortex, ROIs were drawn from the approximate medial border with the Retrosplenial Area (RSPd) to the border with the piriform area (PIR). Cortical barrels within layer 4 (‘L4’) of the somatosensory cortex were used as an anatomical landmark to delineate the ‘upper’ cortex (L1-L4) from the ‘lower’ cortex (L5-L6). Only the upper cortex was taken forward for downstream analysis. For the hippocampus, a single ROI was drawn to encompass the whole hippocampus, excepting the stratum oriens (SO). ROIs for both regions were then isolated and preprocessed via median filter (pixel radius = 1); followed by rolling ball background subtraction (pixel radius = 40). Preprocessed images were thresholded via Maximum Entropy, followed by application of the Watershed algorithm available to download from: <https://github.com/krsholt/FIREPS1_Code_Macros>. Thresholding was performed on the basis of staining batch, and all images within a batch, from a given channel, were thresholded using identical parameters. For burden analysis, the % ROI area occupied by thresholded objects in each channel was quantified using the ‘Measure’ function in FIJI/ImageJ. For counts of absolute object numbers in each channel, the ‘Count Particles’ function was applied.

***Protein extraction and fractionation and western blotting*:** For fractionation of Aβ protein, a procedure similar to that previously described was employed ^9^. The tissue was thawed, weighed and homogenised in Triton-X-lysis buffer containing 1% Triton^TM^ X-100, 50 mM Tris-HCl, pH 7.6, 150 mM NaCl, 2 mM EDTA, 1 mM DTT, PhosSTOP™ phosphatase inhibitor tablet (Roche) and cOmplete™, EDTA-free Protease Inhibitor Cocktail tablet (Roche) at a ratio of 1:10 (1 mg tissue = 10 μL Triton-X-lysis buffer). The resulting total lysates were assayed for protein concentration using the Pierce^TM^ BCA protein assay. Triton-X-soluble, SDS-soluble and urea-soluble fractions were generated for cortical and hippocampal tissue. Equal amounts of Triton-X-lysate protein (200 µg) in a final volume of 100 µL were centrifuged at 100,000 × g for 60 min at 4 °C. The supernatant was collected as the Triton-X-soluble fraction. Pellets were washed with Triton-X-lysis buffer and then re-suspended in equals volumes of Triton-X-lysis buffer containing 1% SDS and centrifuged at 100,000 × g for 60 min at 4 °C to remove SDS-soluble protein. The remaining pellets were washed with Triton-X-lysis buffer containing 1% SDS and dissolved in buffer containing 30 mM Tris-HCl, pH 8.5, 7 M urea, 2 M thiourea and 4% CHAPS and centrifuged at 100,000 × g for 60 min at 4 °C to generate the urea-soluble fraction for analysis by western blot.

***Western blotting:*** Protein samples were separated by gel electrophoresis using a Tricine-based buffer system to ensure maximum resolution of proteins in the molecular weight range of interest. Approximately 5 μg of protein was loaded onto a 10% Tris-Tricine polyacrylamide gel (Invitrogen) and subjected to electrophoresis (120V, 80 min). The gels were blotted onto 0.45μm pore PVDF membranes (Merck) using the Xcell Surelock system (Invitrogen) according to the manufacturer’s instructions. Membranes were then blocked for 1 h at room temperature with 5% (w/v) non-fat dried milk in TBS with 0.1% Tween 20. The membranes were incubated at 4 °C overnight with primary antibodies diluted in blocking solution and visualised using HRP-based secondary antibodies (Anti-rabbit 1:2000, Anti-mouse 1:1000) followed by chemiluminescent detection on Amersham Hyperfilm ECL films. Western blots were analysed by digitally scanning the blots, followed by densitometric analysis (ImageJ) and normalisation against the signal obtained by reprobing the membranes with anti-β-actin. Antibodies, dilutions, techniques, sources, and catalogue/product numbers used are as follows: 6E10 (#803003, 1:1000, Biolegend), APP C-terminus C1/6.1 (#802801, 1:1000 Biolegend), ß-actin (#ab8227, 1:10,000, Abcam).

***Acquisition of plaque images and plaque property analysis:*** Immunofluorescence staining was conducted for plaque properties, utilizing OC (fibrillar amyloid) and Methoxy-X-04 (ß-sheet). For the analysis of neurite dystrophies, Methoxy-X-04 and SMI312 (neurofilament) were used. Leica SP8 confocal microscope equipped with a 63x oil immersion objective was used for image acquisition. When imaging plaques, a single plaque was centered in the field of view, and sequential imaging was performed with 1 μm increments on the z-axis from the top to the bottom of the plaque. For 20x imaging, a z-stack was captured at 2 μm increments over 12 μm, with subsequent maximum projection for 2D analysis. For neurite debris imaging, high intensity staining at the extremes of the stack was excluded to prevent interference with downstream analysis. All imaging was executed blinded to genotype. For image pre-processing, visualization of Leica Image Format (.lif) files – in 3D and 2D - was conducted using Arivis Vision4D v4.0.0. Image processing was completed concurrently using custom-built Python 3.12 scripts written in Jupyter Notebooks, in Visual Studio Code v1.84.0, and run on MacBook Pro M2 Max. Image pixel values were rescaled to between 0-255 intensity values, standardized over the range to improve contrast, and denoised using bilateral or Gaussian denoise algorithms. Particle enhancement was employed for punctate staining. Python packages used: OpenCV, Scikit-Image, and SciPy.NDimage.

For machine learning-based object segmentation, 3D objects were segmented from pre-processed images in Vision4D’s Machine Learning Classifier function. 10-15 image sections, each ~20 pixels in size, were manually drawn per desired object class e.g., plaque compartment or dystrophic neurite. Manually drawn image sections were selected from ~6 images across genotypes per classification. A 3D voxel-based random forest classification approach was then employed, incorporating the voxel microenvironment information from the manually classified segments into the training dataset. Automatic train-test split, hyperparameter tuning, cross-validation, and implementation were performed in the program. The classification results per object class were stored as labelled objects in OME TIFF files and imported into Python 3.12 for downstream analysis.

OME TIFF files containing labelled objects were imported into a Python pipeline for further image handling and post-processing. All segmented objects were first binarized and closed (small objects removed and gaps closed) using SciPy.NDImage. Object classification often resulted in multiple single objects clustering together as one plaque. To combine these objects for downstream analysis, the stack size of plaque images was reduced on the X & Y axes to the central cuboid containing the plaque with ~30µm empty border from surface to image edge. Objects were identified in this reduced 3D space, Euclidean distance clustering was then employed from SciPy.NDImage to merge them into a single, larger plaque object.

Morphological properties of the combined volumes were calculated using parameters built into Scikit-Image ‘regionprops’ function and OpenCV in Python 3.12. Properties calculated were: Volume (V), surface area (SA; Knud Thomsen ellipsoid surface area formula), extent (within the bounding cuboid), complexity (V/SA), sphericity (Ψ = (π^(1/3) * (6 * V)^(2/3)) / SA), solidity (Ratio of pixels in the object region to pixels of the convex hull image), and major and minor axis length (longest and shortest axes). The ratio between the total volume of OC (fibrillar amyloid) and the total volume of Methoxy-X-04 (ß-sheet) was calculated and added to list of morphological features.

Antibody SMI312 was used to stain for healthy and dystrophic neurites proximal and distal to amyloid-ß plaques. Segmented dystrophy objects were binarized and closed using SciPy.NDImage. Dystrophy centroids were quantified either in relation to the plaque surface mesh, in APP genotypes, or in the volume of the total image, in plaque distal and non-APP genotypes. In plaque-proximal conditions, the distance of each dystrophy to the plaque surface was calculated using SciPy Euclidean Distance algorithm, and dystrophies outside a 50μm radius from the plaque surface were filtered out. The number of dystrophies proximal to the plaque was then normalized to the volume of the 50μm surrounding region used in the quantification. For dystrophies distal to the plaque and in non-APP genotypes, individual dystrophy centroids were quantified in and normalized to the volume of the total image.

A machine learning classifier model was employed to predict the genotype (APP-WT, APP-FIRE) from the plaque compartment (OC & MX04) morphological features: volume, extent, surface area, complexity, sphericity, solidity, volume ratio, major and minor axis length. Decision Tree, Random Forest, and Support Vector Machine (SVM) classifiers were compared using Python's Scikit-learn framework. Hyperparameter tuning was performed using grid search, for Decision Tree & SVM, or random search, for Random Forest models. Cross-validation (5-fold) of results was conducted, and the model with the most balanced accuracy, sensitivity, recall, and F1 score was selected. In this case, a Decision Tree was selected for downstream analysis. Extraction of feature scores and visualization of the decision tree were conducted to elucidate the learning process and identify key predictive plaque properties. The model parameters were saved as a .pkl file. All machine learning analysis was conducted in Jupyter Labs on a MacBook Pro M2 Max.

***Array Tomography:*** Tissue samples from the somatosensory cortex were dissected in half to form two segments of approximate 1 mm^3^ dimension. Segments were then processed in accordance with methods described previously ^10^. Briefly, tissue was fixed in 4% paraformaldehyde +2.5% sucrose for 3 hours; dehydrated in ascending ethanol series (50%, 75%, 90%, 100%) and embedded in 100% LR white acrylic resin. Embedded tissue was sectioned into ultrathin ribbons of contiguous 70 nm thick tissue slices. Immunostaining of ribbons was then performed as follows: ribbons were washed 5 times via continuous flow with 1X tris-buffered saline (TBS) and rehydrated with filtered 50 mM glycine solution (0.188g glycine + 50 mL TBS) for 10 minutes at room temperature. Tissue was then washed a further 5 times before blocking for 30 minutes at room temperature using filtered AT blocking reagent (0.05 g gelatin from cold water fish skin + 25 µM TWEEN-20 in 50 mL TBS). Tissue was washed a further 5 times prior to incubation with primary antibodies (anti-synaptophysin [SY38] # ab8049, 1:50 Abcam; anti-PSD95 XP #3450S, 1:50 Cell signalling Technology) overnight at 4^o^C. The next day, primary antibodies were eluted, and tissue washed 5 times before incubation with secondary antibodies (anti-mouse-Cy3 #715-165-150, 1:50 Jackson Immunoresearch; anti-rabbit-Alexa488 # A21206 1:50 Invitrogen; anti-chicken-Alexa647 # A78952 1:50 Invitrogen) for 30 minutes at room temperature while shielded from light. Secondary antibodies were then eluted, and the tissue washed 5 times followed by incubation with 150 µM Methoxy-X04 (Sigma-Aldrich; C5135-500G) for 15 minutes at room temperature, protected from light. A final 5 washes were then performed and ribbons mounted on to 25 x 75 x 1mm Superfrost Plus Adhesion microscope slides (Epredia; J1800AMNZ) using ShandonTM Immu-Mount (Fisher Scientific; FIS9990402).

Imaging of array tomography ribbons was performed via Zeiss Axio Imager Z2 epifluorescent microscope, equipped with a CoolSnap digital camera (Zeiss, Baden-Württemberg, Germany). Image acquisition was carried out using a 63X 1.4 NA Plan Apochromat objective. Serial images were produced along the length of each ribbon by identifying a landmark that tracked through contiguous sections. Raw Images were acquired using ZEN Blue 3.0 software and were saved in raw 16-bit .czi format (1392x1040p) Image series were stacked into 3-dimensional volumes using FIJI (ImageJ) using macros available at the previously linked online repository. Stacks were then aligned via rigid and affine registration, segmented by staining batch and data extracted using proprietary ImageJ, MATLAB and Python scripts, all of which are available freely at: [*https://github.com/Spires-Jones-Lab*](https://github.com/Spires-Jones-Lab). All images from a given channel were segmented using the same parameters by an experimenter blind to genotype. For analysis of synapse density in the neuropil, synaptophysin (SY38) was used as the reference for generating neuropil masks for each image stack. For analysis of object colocalisation, a minimum distance of 0.5 microns between object centroids was used as the parameter for defining pre- and post-synaptic pairing (“paired synapses”).

***Data availability statement:*** Accession code relating to RNA-seq data (ArrayExpress) are E-MTAB-14231 and E-MTAB-14233. Data that support the findings of this study are either either available in the paper and its supplementary information or available from the authors on request.

***Statistics:*** Throughout the manuscript the statistical test is stated, along with p values, and test statistic. Statistical tests performed are two-sided, and performed on biological replicates, not technical replicates. Differential expression analysis of RNA-seq data was performed using DESeq2 88, with a significance threshold calculated at a Benjamini–Hochberg-adjusted P value of <0.05. For cell counting and the experimenter was blind to the condition or genotype. For all images displayed, contrast and brightness are applied in a linear manner across the entire image.

**References**

1	Rojo, R. *et al.* Deletion of a Csf1r enhancer selectively impacts CSF1R expression and development of tissue macrophage populations. *Nat Commun* **10**, 3215 (2019). <https://doi.org/10.1038/s41467-019-11053-8>

2	Jankowsky, J. L. *et al.* Mutant presenilins specifically elevate the levels of the 42 residue beta-amyloid peptide in vivo: evidence for augmentation of a 42-specific gamma secretase. *Human molecular genetics* **13**, 159-170 (2004). <https://doi.org/10.1093/hmg/ddh019>

3	Zheng, G. X. *et al.* Massively parallel digital transcriptional profiling of single cells. *Nat Commun* **8**, 14049 (2017). <https://doi.org/10.1038/ncomms14049>

4	Hao, Y. *et al.* Integrated analysis of multimodal single-cell data. *Cell* **184**, 3573-3587 e3529 (2021). <https://doi.org/10.1016/j.cell.2021.04.048>

5	Germain, P. L., Lun, A., Garcia Meixide, C., Macnair, W. & Robinson, M. D. Doublet identification in single-cell sequencing data using scDblFinder. *F1000Res* **10**, 979 (2021). <https://doi.org/10.12688/f1000research.73600.2>

6	Dobin, A. *et al.* STAR: ultrafast universal RNA-seq aligner. *Bioinformatics* **29**, 15-21 (2013). <https://doi.org/bts635> [pii]

10.1093/bioinformatics/bts635

7	Liao, Y., Smyth, G. K. & Shi, W. featureCounts: an efficient general purpose program for assigning sequence reads to genomic features. *Bioinformatics* **30**, 923-930 (2014). <https://doi.org/btt656> [pii]

10.1093/bioinformatics/btt656

8	Love, M. I., Huber, W. & Anders, S. Moderated estimation of fold change and dispersion for RNA-seq data with DESeq2. *Genome Biol* **15**, 550 (2014). <https://doi.org/s13059-014-0550-8> [pii]

10.1186/s13059-014-0550-8

9	Jiwaji, Z. *et al.* Reactive astrocytes acquire neuroprotective as well as deleterious signatures in response to Tau and Aß pathology. *Nat Commun* **13**, 135 (2022). <https://doi.org/10.1038/s41467-021-27702-w>

10	Kay, K. R. *et al.* Studying synapses in human brain with array tomography and electron microscopy. *Nat Protoc* **8**, 1366-1380 (2013). <https://doi.org/nprot.2013.078> [pii]

10.1038/nprot.2013.078
